# Adaptation of prokaryotic toxins for negative selection and cloning-independent markerless mutagenesis (CIMM) in *Streptococcus* species

**DOI:** 10.1101/2023.01.03.522674

**Authors:** Lena Li, Hua Qin, Zhengzhong Zou, Jens Kreth, Justin Merritt

## Abstract

The *Streptococcus mutans* genetic system offers a variety of strategies to rapidly engineer targeted chromosomal mutations. Previously, we reported the first *S. mutans* negative selection system that functions in a wild-type background. This system utilizes induced sensitivity to the toxic amino acid analog *p-*chlorophenylalanine (4-CP) as a negative selection mechanism, and was developed for counterselection-based cloning-independent markerless mutagenesis (CIMM). While we have employed this system extensively for our ongoing genetic studies, we have encountered a couple limitations with the system, mainly its narrow host range and the requirement for selection on a toxic substrate. Here, we report the development of a new negative selection system that addresses both limitations, while still retaining the utility of the previous 4-CP-based markerless mutagenesis system. We placed a variety of toxin-encoding genes under the control of the xylose-inducible Xyl-S expression cassette and found the Fst-sm and ParE toxins to be suitable candidates for inducible negative selection. We combined the inducible toxins with an antibiotic resistance gene to create several different counterselection cassettes. The most broadly useful of these contained a wild-type *fst-sm* open reading frame transcriptionally fused to a point mutant form of the Xyl-S expression system, which we subsequently named as IFDC4. IFDC4 was shown to exhibit exceptionally low background resistance, with 3 – 4 log reductions in cell number observed when plating on xylose-supplemented media. IFDC4 also functioned similarly in multiple strains of *S. mutans* as well as with *S. gordonii* and *S. sanguinis*. We performed CIMM with IFDC4 and successfully engineered a variety of different types of markerless mutations in all three species. The counterselection strategy described here provides a template approach that should be adaptable for the creation of similar counterselection systems in many other bacteria.

## Introduction

Many *Streptococcus* species are highly amenable to genetic manipulation as a consequence of their efficient natural competence machinery (1, 2). Accordingly, a variety of sophisticated genetic systems have also been developed for molecular pathogenesis studies of certain medically significant *Streptococcus* species, such as the prominent oral pathobiont *Streptococcus mutans* (3–6). The *S. mutans* genetic system has now evolved to the point where targeted chromosomal mutations can be reliably engineered within a matter of days using various cloning-independent mutagenesis strategies. Such approaches circumvent the requirement for intermediate hosts like *E. coli* during construct assembly, which significantly reduces the time and effort required to engineer mutations of interest. The vast majority of targeted mutations in *S. mutans* (and other streptococci) are created using marked mutagenesis, typically as allelic replacements with antibiotic resistance cassettes. While simple to engineer, marked mutations have a couple fundamental limitations that can be highly problematic for genetic studies. Firstly, antibiotic resistance cassettes normally contain promoters and/or transcription terminators, which can introduce significant polar effects altering the expression patterns of genes downstream of a mutation site (7). Secondly, the number of individual mutations that can be engineered into a single strain is inherently limited by the number of unique selectable markers available for use in a particular organism. Both of these limitations can by addressed through the creation of markerless mutations, but unfortunately, only a limited number of organisms have markerless mutagenesis systems available for use.

Most markerless mutations are created by first employing marked mutagenesis to insert an antibiotic resistance cassette onto a chromosome followed by a second step to subsequently remove the cassette to create the final markerless mutant strain. Once an antibiotic resistance cassette has been removed, the same procedure can be continually repeated to engineer any number of additional markerless mutations in the same strain (7). There are several common strategies employed to remove antibiotic resistance cassettes from the chromosome and all have been previously employed in *S. mutans*. The first is a two-step integration and excision strategy using a conditionally replicating temperature sensitive plasmid (8). Temperature-sensitive plasmids are only available for a limited number of organisms, and their temperature sensitivity often lacks stringency. The next strategy uses site-specific recombinases, such as Cre/LoxP to remove antibiotic resistance cassettes following their insertion (9). This approach has the major advantage of being theoretically adaptable for a wide range of species, but also typically relies upon temperature sensitive vectors to provide transient *cre* expression. Cre-mediated recombination between LoxP sites also generates scars on the chromosome that can interfere with the creation of certain types of mutations like point mutations or specific gene truncations (10). The third common strategy for markerless mutagenesis utilizes a combination of both positive and negative selection (i.e. counterselection). When available for use, the counterselection approach is often preferred due to its ease of use and suitability for the creation of all types of mutations, such as deletions, insertions, truncations, point mutations, etc. (7, 11). The principal limitation is that few efficacious negative selection markers are available for use in most bacteria.

We previously developed the first counterselection-based markerless mutagenesis system available for *S. mutans*, using induced sensitivity to galactose as a negative selection mechanism (12). By creating a mutant recipient strain defective in galactose catabolism, it was possible to employ the endogenous *S. mutans galK* gene (encoding galactokinase) as a negative selection marker in the presence of galactose-supplemented media. The obvious drawback is that this approach is not compatible with wild-type strains of *S. mutans* or any of the numerous other bacterial species that naturally catabolize galactose. This issue has limited the widespread adoption of the *galK* negative selection system. To improve upon this major shortcoming, we next adapted the PheS/*p*-chlorophenylalanine (4-CP) negative selection system for use in *S. mutans* markerless mutagenesis (7, 13, 14). For this approach, an A314G mutant version of the endogenous *S. mutans* PheS protein is expressed for negative selection on media supplemented with the toxic phenylalanine analog 4-CP. *S. mutans* strains producing the A314G mutant PheS protein exhibit much higher sensitivity to 4-CP toxicity compared to the wildtype, thus facilitating negative selection. The major advantage to this approach is that the *pheS* gene is highly conserved among prokaryotes, and consequently, 4-CP negative selection is theoretically adaptable for use in most wild-type bacteria, unlike other negative selection approaches (7). To create a counterselection cassette, we combined an antibiotic resistance gene and a point mutant *S. mutans pheS* into a single cassette referred to as IFDC2. We further illustrated how to employ the IFDC2 cassette for the first demonstration of cloning-independent markerless mutagenesis (CIMM) (7). Despite the major improvements of the CIMM approach over previous markerless mutagenesis strategies, several limitations were still observed. Background 4-CP resistance was often higher than desired and the IFDC2 cassette only functioned in *S. mutans*. In our experience, reliable 4-CP negative selection requires mutant *pheS* genes to be derived from the organism of interest. In a subsequent study, we were able to further increase the stringency of 4-CP negative selection in *S. mutans* by adding a second point mutation to the *pheS* gene contained on IFDC2, resulting in a T260S PheS mutation in addition to the prior A314G mutation (15). However, this updated version (referred to as IFDC3) still retained the requirement for species-specific *pheS* genes to be employed. In addition, while using 4-CP-based negative selection, we have encountered instances in which a particular chromosomal mutation sensitized the strain to 4-CP, thus rendering the markerless mutant difficult to isolate on 4-CP plates. For this reason, we were interested to search for an alternative negative selection mechanism that does not require plating on media supplemented with a toxic substrate like 4-CP. Additionally, we were interested to develop a single counterselection cassette that functions in multiple streptococci, unlike our *pheS-*based systems. Here, we describe the development of a toxin-based counterselection mechanism that satisfies these requirements, while also exhibiting exceptionally low background resistance.

## Materials and Methods

### Primers, bacterial strains, and growth conditions

The primers and bacterial strains used in this study are shown in Tables S1 and S2, respectively. *S. mutans* strains UA159, UA140, JF243, CL1, and their derivatives were cultured anaerobically (in an atmosphere consisting of 85% N_2_, 10% CO_2_, and 5% H_2_) at 37 °C in Todd-Hewitt medium (Difco) supplemented with 0.3% (wt vol^−1^) yeast extract (THYE) or on THYE agar plates. For the selection of antibiotic resistant colonies, 1 mg mL^−1^ spectinomycin (Sigma-Aldrich) was added to growth media. *Streptococcus sanguinis* strain SK36, *Streptococcus gordonii* strain DL1, and their derivatives were cultured in 5% CO_2_ at 37 °C in Brain Heart Infusion medium (BHI; Difco) or on BHI agar plates. For the selection of antibiotic resistant colonies, 400 μg mL^−1^ spectinomycin (Sigma-Aldrich) was added to growth media.

### Transformation

DNA constructs were introduced into *S. mutans* using a previously described methodology (2). Briefly, *S. mutans* cultures were diluted 1:40 from overnight cultures and grown to an optical density of OD_600_ ∼0.1 in THYE before the addition of transforming DNA and 1 μg ml^−1^ Competence Stimulating Peptide (CSP; GenScript). The cultures were subsequently incubated for an additional 2 h and then plated on antibiotic-supplemented THYE plates. Transformation of *Streptococcus sanguinis* and *Streptococcus gordonii* were performed as described previously (16). Briefly, cultures were diluted 1:40 from overnight cultures and grown to an optical density of OD_600_ ∼0.07 in BHI medium before the addition of transforming DNA and 1 μg ml^−1^ of the appropriate CSP (ChemPep). The cultures were subsequently incubated for an additional 2 h and then plated on antibiotic-supplemented BHI plates. For strains undergoing negative selection, THYE or BHI plates were supplemented with 1% (wt vol^−1^) xylose (Sigma-Aldrich).

### DNA manipulation

Phusion DNA polymerase (Thermo Scientific) or AccuPrime Polymerase (Invitrogen) were used to amplify individual PCR amplicons and overlap extension PCR (OE-PCR) products.

### Generation of strains harboring candidate counterselection cassettes

Each candidate counterselection cassette was created using OE-PCR to transcriptionally fuse a toxic gene product to the Xyl-S xylose induction cassette (17) for negative selection and then it was subsequently combined with a downstream spectinomycin resistance gene *aad9* for positive selection. Each of the candidate counterselection cassettes was inserted into the *brsRM* locus of *S. mutans* (18) to assay its functionality. The Fst-sm-encoding counterselection cassette was the first to be constructed. The Xyl-S induction cassette and spectinomycin resistance gene *aad9* were both PCR amplified from the plasmid pZX9 (17) using the primer pairs 1925-2 Fwd + 1925-2 Rvs and 1925-4 Fwd + 1925-IFDC-Rvs, respectively. The *fst-sm* gene was PCR amplified using UA159 genomic DNA (gDNA) and the primer pair 1925-3 Fst-sm Fwd + 1925-3 Fst-sm Rvs. Each of the resulting PCR amplicons contain segments of sequence complementarity facilitating their subsequently assembly via OE-PCR with the primer pair 1925-2 Fwd + 1925-IFDC-Rvs. Next, the upstream and downstream homologous fragments used for targeting recombination within the *brsRM* locus were PCR amplified from UA159 gDNA using the primer pairs 1925-1 brsRM159-LF + 1925-1 Rvs and 2018-2 Fwd + 1925-5 brsRM159-RR, respectively. The resulting PCR amplicons were mixed with that of the previously assembled inducible *fst-sm* construct and assembled into a final construct via OE-PCR using the primer pair 1925-1 brsRM159-LF + 1925-5 brsRM159-RR. The resulting full-length construct was then transformed to UA159 and selected on antibiotic-supplemented agar plates. Several clones of the resulting transformation were tested for negative selection in the presence of xylose. Clones of interest were sequenced to confirm the expected counterselection cassette genotype. Functional counterselection cassettes contained either of two point mutations located within codon 7 of the *xylR* open reading frame (ORF), resulting in either an A7S or A7T mutation within XylR. These two strains were subsequently named as 159MfstS (XylR A7S) and 159MfstT (XylR A7T). To generate counterselection cassettes encoding MazF, SmuT, and RNaseH, similar assembly strategies were performed as described for strains 159MfstS and 159MfstT, except that the following primers pairs were employed. The *mazF* construct utilized primers 1925-2 Fwd + 1915-2M Rvs, 1915-3 MazF Fwd + 1915-3 MazF Rvs, and 1915-4M Fwd + 1925-IFDC-Rvs to generate the strain 159MmazF. The *smuT* construct utilized primers 1925-2 Fwd + 1915-2S Rvs, 1915-3 SmuT Fwd + 1915-3 SmuT Rvs, and 1915-4S Fwd + 1925-IFDC-Rvs to generate the strain 159MsmuT.

The *rnH* construct utilized primers 1925-2 Fwd + 1917-2R Rvs, 1917-3 RnaseH Fwd + 1917-3 RnaseH Rvs, and 1917-4R Fwd + 1925-IFDC-Rvs to generate the strain 159MrnH. To create a counterselection cassette containing the *parE* toxin, strain 159MfstS was used as a template to amplify the upstream homologous fragment and Xyl-S induction cassette using the primers 1925-1 brsRM159-LF + 2019-1-Rvs. The spectinomycin resistance gene *aad9* and the downstream homologous fragment were similarly amplified from strain 159MfstS using the primers 2019-3 Fwd + 1925-5 brsRM159-RR. The *parE* ORF was PCR amplified with the primers 2019-2 ORF5-Fwd + 2019-2 ORF5-Rvs together with lysates of *S. mutans* strain UA140, which naturally harbors the cryptic plasmid pUA140 containing the *parE* addiction module. Each of these PCR amplicons was mixed and assembled via OE-PCR using the primers 1925-1 brsRM159-LF + 1925-5 brsRM159-RR. The assembled construct was subsequently transformed to UA159 and selected on antibiotic-supplemented agar plates. Several clones of the resulting transformation were tested for negative selection in the presence of xylose. Clones of interest were sequenced to confirm the expected counterselection cassette genotype. Functional counterselection cassettes were found to contain one of two point mutations located either within the *parE* ribosome binding site (RBS) or within the *parE* ORF, encoding a ParE S56G mutation. These two strains were subsequently named 159MparE1 (ParE RBS) and 159MparE2 (ParE S56G). To compare the functionality of the *fst-sm-* and *parE*-containing counterselection cassettes in other *S. mutans* strains, the cassettes were first PCR amplified from strains 159MfstS, 159MfstT, 159MparE1, and 159MparE2 using the primers 1925-1 brsRM159-LF + 1925-5 brsRM159-RR. These amplicons were subsequently transformed into *S. mutans* wild-type strains CL1, JF243, and UA140 and selected on antibiotic-supplemented agar plates. The resulting strains were named CL1MfstS, CL1MfstT, JF243MfstS, JF243MfstT, 140MfstS, and 140MfstT (Fst-sm) as well as CL1MparE1, CL1MparE2, JF243MparE1, JF243MparE2, 140MparE1, and 140MparE2 (ParE).

To test the functionality of the counterselection cassettes in both *Streptococcus sanguinis* and *Streptococcus gordonii*, candidate cassettes were used to create allelic replacement mutants of the *spxB* genes from both species. For *S. sanguinis*, the *spxB* upstream and downstream homologous fragments were PCR amplified from wild-type strain SK36 using the primer pairs 2010-1-LF + 2110-1-LR and 2110-3-RF + 2110-3-RR, respectively. The counterselection cassettes were PCR amplified from *S. mutans* strains 159MfstS, 159MfstT, 159MparE1, and 159MparE2 using the primers 2110-2 ΔspxB Fwd + 2110-2 ΔspxB Rvs. The upstream and downstream homologous fragments were mixed with each of the counterselection cassette amplicons and assembled via OE-PCR using the primers 2010-1-LF + 2110-3-RR. These amplicons were subsequently transformed into *S. sanguinis* strain SK36 and selected on antibiotic-supplemented agar plates. The resulting strains were named SK36SBfstS and SK36SBfstT (Fst-sm) as well as SK36SBparE1 and SK36SBparE2 (ParE). For *S. gordonii*, the *spxB* upstream and downstream homologous fragments were PCR amplified from wild-type strain DL1 using the primer pairs 2021-22-1-LF + 2021-22-1-ΔspxB-Rvs and 2021-22-3-ΔspxB-Fwd + 2021-22-3-RR, respectively. The counterselection cassettes were PCR amplified from *S. mutans* strains 159MfstS, 159MfstT, 159MparE1, and 159MparE2 using the primers 1925-IFDC-Fwd + 1925-IFDC-Rvs. The upstream and downstream homologous fragments were mixed with each of the counterselection cassette amplicons and assembled via OE-PCR using the primers 2021-22-4 nest Fwd + 2021-22-4 nest Rvs. These amplicons were subsequently transformed into *S. gordonii* strain DL1 and selected on antibiotic-supplemented agar plates. The resulting strains were named DL1SBfstS and DL1SBfstT (Fst-sm) as well as DL1SBparE1 and DL1SBparE2 (ParE).

### Markerless replacement of *brsM* with the β-glucoronidase-encoding gene *gusA*

In order to check the efficiency of markerless allelic replacements, gDNA from strain 159MGusA was used a template for PCR using the primer pair 1925-1 brsRM159-LF + 1925-5 brsRM159-RR. This PCR amplicon was transformed into strains 159MfstS, 159MfstT, 159MparE1, and 159MparE2 to replace the *brsM* open reading frame (ORF) with the *gusA* ORF and remove the respective counterselection cassettes. Transformants were selected on THYE plates supplemented with 1% (wt vol^−1^) xylose and 200 (μg ml^−1^) x-Gluc. The generation of the expected Δ*brsM gusA*^+^ genotype results in transformants with blue color, while background growth of the parent strain results in white colonies.

### Creation of markerless *spxB-renG* green renilla luciferase reporters in *S. sanguinis* and *S. gordonii*

To create the markerless *spxB-renG* transcription fusion in *S. sanguinis* strain SK36, we transformed the *renG* construct into a previously constructed *fst-sm* counterselection cassette allelic replacement of the *spxB* ORF. The SK36 *spxB* upstream homologous fragment was PCR amplified from SK36 gDNA using the primer pair 2110-1-LF + 2021-17-1-Rvs, while the downstream homologous fragment was PCR amplified using the same gDNA template with the primer pair 2021-17-3-Fwd + 2021-17-3-Rvs. The *renG* ORF was PCR amplified from the gDNA of a previously constructed *comX*-*renG* reporter strain of *S. mutans* using the primer pair 2021-17-2 Fwd renG + 2021-17-2 Rvs renG. The resulting PCR amplicons were mixed and assembled via OE-PCR using the primer pair 2110-1-LF + 2021-17-3-Rvs. The OE-PCR amplicon was subsequently transformed into strain SK36SBfstT and selected on BHI agar plates supplemented with 1% (wt vol^−1^) xylose to generate the markerless *spxB-renG* reporter strain SK36RGfstT.

To create the markerless *spxB-renG* transcription fusion in *S. gordonii* strain DL1, we transformed the *renG* construct into a previously constructed *fst-sm* counterselection cassette allelic replacement of the *spxB* ORF. The DL1 *spxB* upstream homologous fragment was PCR amplified from DL1 gDNA using the primer pair 2021-22-1-LF + 2021-23-1-Rvs, while the downstream homologous fragment was PCR amplified using the same gDNA template with the primer pair 2021-23-3-Fwd + 2021-23-3-Rvs. The *renG* ORF was PCR amplified from the gDNA of a previously constructed *comX*-*renG* reporter strain of *S. mutans* using the primer pair 2021-17-2 Fwd renG + 2021-17-2 Rvs renG. The resulting PCR amplicons were mixed and assembled via OE-PCR using the primer pair 2021-22-1-LF + 2021-23-3-Rvs. The OE-PCR amplicon was subsequently transformed into strain DL1SBfstT and selected on BHI agar plates supplemented with 1% (wt vol^−1^) xylose to generate the markerless *spxB-renG* reporter strain DL1RGfstT.

### Generation of markerless *galK* nonsense point mutations in *S. mutans* and *S. gordonii*

To engineer a markerless point mutation into the *S. mutans galK* gene, we first inserted the *fst-sm* (A7T) counterselection cassette between the *galR* and *galK* ORFs on the chromosome of *S. mutans* strain UA159. The upstream and downstream homologous fragments were PCR amplified from UA159 using the primer pairs 2130-1Fwd LAM + 2130-1Rvs LAM and 2130-3Fwd RAM + 2130-3 Rvs RAM, respectively. The *fst-sm* (A7T) counterselection cassette was amplified from *S. mutans* strain 159MfstT using the primer pair 1925-IFDC-Fwd + 1925-IFDC-Rvs. The three PCR amplicons were mixed and subsequently assembled using OE-PCR with the primer pair 2130-4 nest Fwd + 2130-4 nest Rvs. The assembled construct was then transformed to UA159 and selected on antibiotic supplemented agar plates to generate the intermediate strain 159GKfstT. A second construct was created to replace the counterselection cassette with a mutagenized version of the *galK* ORF in which codon 8 was changed from CAA to TAA. The upstream and downstream homologous fragments were PCR amplified from UA159 using the primer pairs 2130-1 Fwd LAM + 2130-5 Rvs galk mut and 2130-6 Fwd galk mut + 2130-3 Rvs RAM, respectively. The PCR amplicons were mixed and assembled via OE-PCR using the primer pair 2130-1Fwd LAM + 2130-3 Rvs RAM to generate the final construct, which was transformed into strain 159GKfstT and selected on xylose-supplement agar plates to generate the strain 159galk22. Several transformants were sequenced to confirm the expected point mutant genotypes.

To generate a markerless *galK* point mutation in *S. gordonii* strain DL1, a similar strategy was employed as described for strain 159galk22. We first inserted the *fst-sm* (A7T) counterselection cassette between the *galR* and *galK* ORFs on the chromosome of DL1. The upstream and downstream homologous fragments were PCR amplified from DL1 using the primer pairs 2201-1 Fwd LAM + 2201-1 Rvs LAM and 2201-3 Fwd RAM + 2201-3 Rvs RAM, respectively. The *fst-sm* (A7T) counterselection cassette was PCR amplified from *S. mutans* strain 159MfstTusing the primer pair 1925-IFDC-Fwd + 1925-IFDC-Rvs. The three PCR amplicons were mixed and subsequently assembled using OE-PCR with the primer pair 2201-4 nest Fwd + 2201-4 nest Rvs. The assembled construct was then transformed to DL1 and selected on antibiotic supplemented agar plates to generate the intermediate strain DL1GKfstT. A second construct was created to replace the counterselection cassette with a mutagenized version of the *galK* ORF in which codon 9 was changed from CAA to TAA. The upstream and downstream homologous fragments were PCR amplified from DL1 using the primer pairs 2201-1 Fwd LAM + 2201-5 Rvs galk mut and 2201-6 Fwd galk mut + 2201-3 Rvs RAM, respectively. The PCR amplicons were mixed and assembled via OE-PCR using the primer pair 2201-1 Fwd LAM + 2201-3 Rvs RAM to generate the final construct, which was transformed into intermediate strain DL1GKfstT and selected on xylose-supplement agar plates to generate the final strain DL1galk25. Several transformants were sequenced to confirm the expected point mutant genotypes.

### Detection of hydrogen peroxide production on Prussian blue indicator plates

The preparation of H_2_O_2_ indicator plates and detection of bacterial H_2_O_2_ production were performed using a previously described methodology (19). Briefly, overnight cultures of each strain were washed twice with PBS and then adjusted to an optical density OD_600_ 0.5. A volume of 10 μl was pipetted onto BHI Prussian-blue indicator plates and incubated overnight at 37 °C in a 5% CO_2_ atmosphere.

### Luciferase assays

Luciferase assays were performed using a previously described methodology (16, 20). Briefly, strains were diluted 1:40 from overnight liquid cultures and then incubated for 4 h at 37 °C. Optical densities (OD_600_) were subsequently measured for each sample to normalize luciferase values. Coelenterazine-h solution (Prolume) was added to each sample at a final concentration of 7.5 μg ml^−1^ immediately followed by measuring the resulting luciferase activity using a Promega Glomax Discover luminometer. Normalized luciferase activity was determined by dividing luciferase relative light units (RLU) by the measured optical density values (RLU/OD_600_).

### Measurement of deoxygalactose sensitivity

Overnight cultures were washed twice with PBS and adjusted to an optical density OD_600_ 0.5. The cultures were serially diluted 10-fold and 10 μl of each dilution was pipetted onto BHI agar plates ± 1% (wt vol^−1^) deoxygalactose. The plates were incubated anaerobically at 37 °C for 24 h.

## Results

### Development of a toxin-based negative selection system in *S. mutans*

Despite the proven utility of 4-CP-based counterselection in *S. mutans*, there are still a couple limitations with this approach that can be problematic, mainly its requirement for a toxic substrate (i.e. 4-CP) and a narrow host range for *pheS*-containing counterselection cassettes. Here, we aimed to improve upon these by developing a new negative selection strategy to facilitate cloning-independent markerless mutagenesis (CIMM). We previously developed a xylose-based gene induction system (called Xyl-S) that was shown to exhibit an exceptionally wide dynamic range with low basal expression, and it also functioned similarly in several different oral streptococci with no evidence of toxicity (17). In fact, we had even employed Xyl-S to engineer multiple conditional lethal mutations in *S. mutans* (17). Thus, we reasoned that the Xyl-S system may also be appropriate to control the expression of a toxic gene product to confer negative selection in *S. mutans* (Fig. 1). As shown in Table 1, we selected several candidate chromosomal toxins from verified *S. mutans* toxin-antitoxin modules (*mazF, smuT*, and *fst-sm*) as well as the plasmid addiction module toxin *parE* from the *S. mutans* cryptic plasmid pUA140 and RNase H (*rnH*) from *E. coli*. Each of these genes was transcriptionally fused to the Xyl-S cassette and transformed into *S. mutans* strain UA159 to test for xylose-inducible negative selection. Following transformation of the constructs, we noted that most seemed to exhibit high levels of basal uninduced toxicity, as some transformants grew slowly and we observed fewer transformants than expected. Regardless, we selected candidate transformants from each reaction and then quantified their viability on agar plates ± 1% (wt vol^−1^) xylose. As shown in Figure 2A, only the *fst-sm* and *parE* constructs inhibited cell growth in the presence of xylose, with each exhibiting potent negative selection. We sequenced several clones of the *fst-sm* and *parE* strains that exhibited inducible negative selection and found all to contain at least one point mutation. For the strains harboring the *fst-sm* construct, they each contained either of two missense mutations within the same codon of the xylose repressor gene *xylR*, conferring either a XylR A7S or A7T mutation (Fig. 2B). The strains harboring *parE* constructs all contained the same XylR A7S mutation because one of the mutated *fst-sm* clones had been employed as a PCR template during the assembly of the *parE* construct (Fig. 2C). However, the *parE* strains also contained one of two additional point mutations, either within the *parE* ribosome binding site (RBS) or within the *parE* ORF, conferring a ParE S56G missense mutation (Fig. 2C). While the impact of the ParE S56G mutation was unclear, the *parE* RBS mutation almost certainly reduced the efficiency of *parE* translation, as the original construct contained a consensus Shine-Dalgarno sequence. Interestingly, we reassembled the *parE* construct with a wild-type *xylR* ORF together with either of the newly identified *parE* RBS or S56G mutations and found that the point mutant *xylR* was indeed required for construct stability. Thus, it would appear that the XylR A7S (and presumably A7T) mutations are responsible for reducing the basal uninduced expression of *fst-sm* and *parE*. The additional compensatory point mutations required for the proper functionality of the *parE* construct suggests that *S. mutans* strains encoding the ParE toxin are likely to be even more sensitive to toxic leaky gene expression as compared to *fst-sm*.

**Table 1.** Candidate toxins examined for inducible negative selection.

| Endogenous Toxin | Source | Citation | Exogenous Toxin | Source | Citation |
| --- | --- | --- | --- | --- | --- |
| **MazF** | TA module on the *S. mutans* chromosome; arrests cell growth via RNA cleavage | (26) | **ParE** | Addiction module of the cryptic *S. mutans* plasmid pUA140; bactericidal via inhibition of DNA gyrase | (27) |
| **SmuT** | TA module on the *S. mutans* chromosome; bactericidal via membrane permeabilization | (28) | **RNase H** | Housekeeping enzyme on the *E. coli* chromosome; arrests cell growth via RNA cleavage | (29) |
| **Fst-sm** | TA module on the *S. mutans* chromosome; putative membrane permeabilization and/or inhibition of cell division | (30) |  |  |  |

**Fig. 1.**
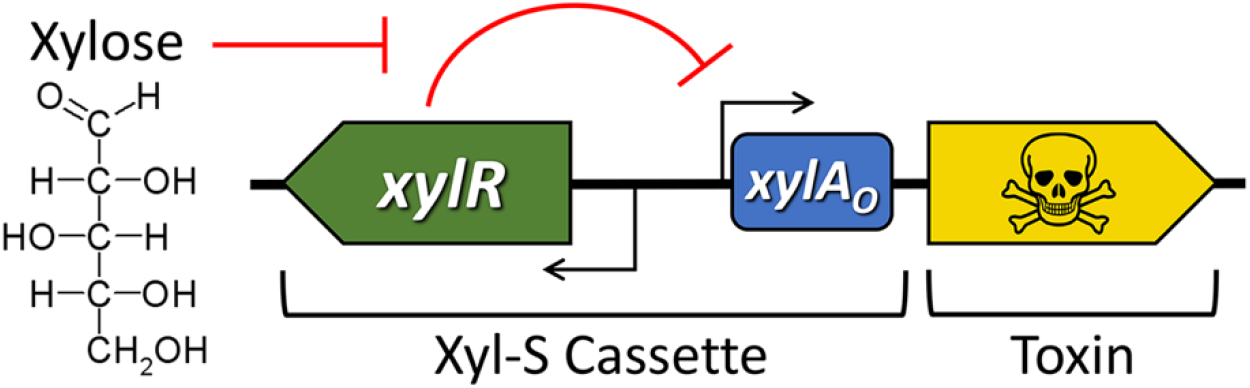
Illustration of the counterselection approach. Putative toxin ORFs (illustrated in yellow) are transcriptionally fused to the xylose-inducible Xyl-S cassette. Xylose is a nontoxic sugar that is not metabolized by streptococci, but is efficiently transported into the cells via unknown mechanisms. Xylose will bind to the xylose repressor XylR and prevent it from inhibiting target gene expression via the xylose operator *xylA*_*O*_.

**Fig. 2.**
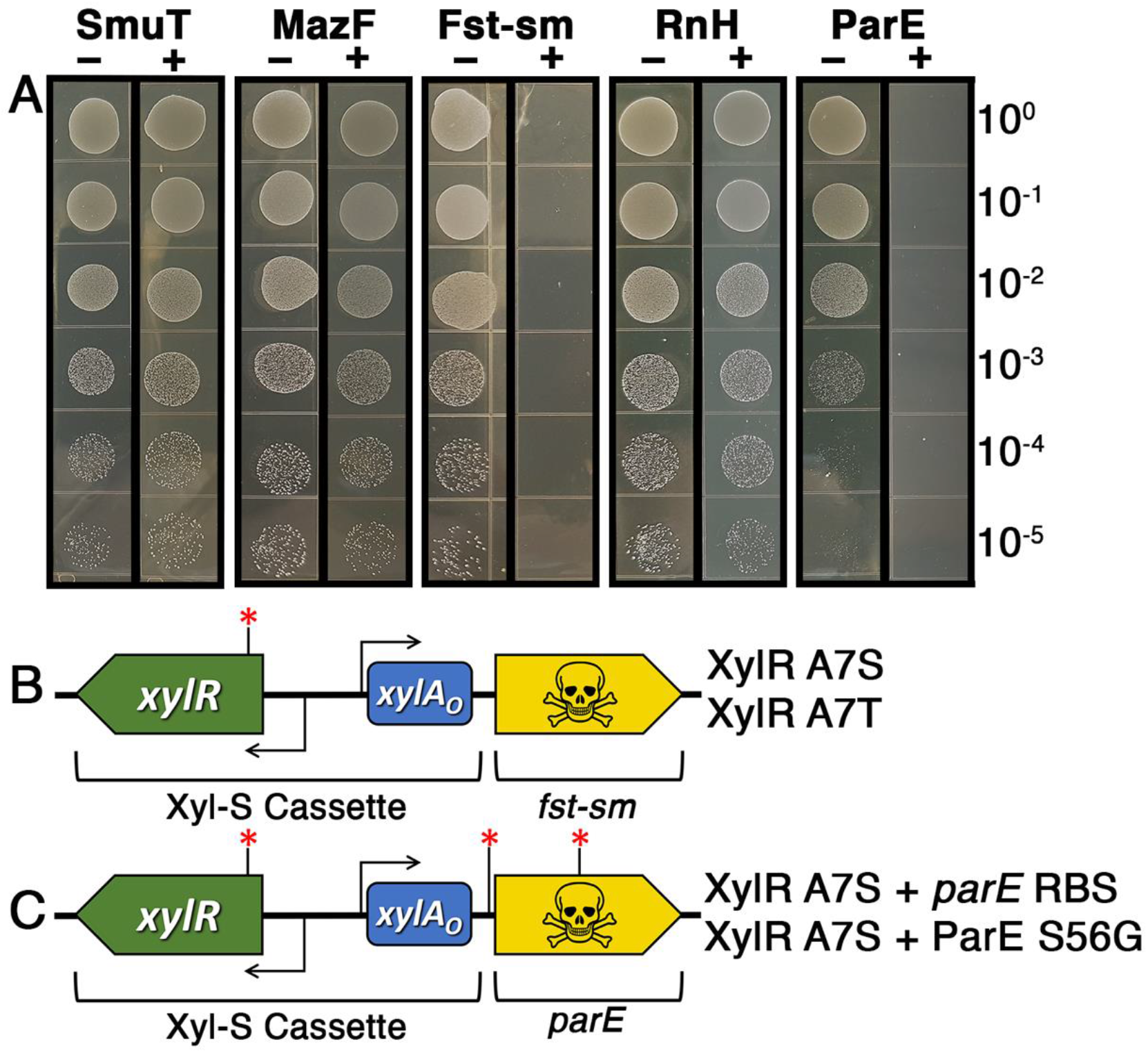
Xylose-inducible negative selection using different toxin genes. A) A mixture of both endogenous and exogenous toxic ORFs were transcriptionally fused to the Xyl-S cassette and then transformed into *S. mutans* strain UA159. The resulting strains were tested for growth on agar plates ± xylose. B) Several clones harboring the *fst-sm* counterselection cassette were sequenced and found to contain one of two different missense mutations within the *xylR* ORF. Both mutations targeted codon 7 of *xylR*, resulting in either A7S or A7T substitutions in XylR. C) Several clones harboring the *parE* counterselection cassette were sequenced and found to contain one of two different point mutations in addition to the same XylR A7S mutation found in some of the *fst-sm* counterselection cassettes. The two unique *parE* mutations occurred either within the *parE* ribosome binding site (RBS) or codon 56 of *parE*, conferring a ParE S56G amino acid substitution.

### Cloning-independent markerless mutagenesis (CIMM) in *S. mutans*

Based upon the extreme potency of the negative selection observed from the *fst-sm* and *parE* counterselection cassettes (Fig. 2A), we were next curious to test the utility of these cassettes for CIMM. As a simple phenotypic readout, we employed the *fst-sm* and *parE* cassettes to engineer markerless *gusA* (β-glucuronidase) replacements of the *S. mutans brsM* ORF. Under normal growth conditions, the gene product of *brsM* inhibits the expression of the *brsRM* operon by preventing BrsR positive feedback autoregulation, leading to extremely low basal expression in the wild-type (Fig. 3A) (18, 21, 22). Consequently, insertion of the *gusA* ORF at the 3’ end of the *brsRM* operon results a white colony phenotype (i.e. low basal *gusA* expression) because the *brsRM* operon remains weakly expressed similar to the parent wild-type (Fig. 3A). Conversely, a *gusA* replacement of *brsM* will stimulate BrsR positive feedback autoregulation of the operon, resulting in strong *gusA* expression and a corresponding blue colony phenotype (Fig. 3A). Using an existing *brsRM-gusA* reporter strain, we first replaced the *gusA* ORF downstream of *brsM* with the *fst-sm* and *parE* counterselection cassettes and subsequently performed a second transformation to replace both *brsM* and the counterselection cassette with *gusA*. As expected, the original parent *brsRM-gusA* reporter strain grew normally on the xylose plates with undetectable β-glucuronidase activity (Fig. 3B). In agreement with the strong xylose-inducible negative selection previously observed with the *fst-sm* and *parE* counterselection cassettes (Fig. 2A), we observed minimal background growth from the negative control transformation reactions (Fig. 3B). In contrast, strains transformed with *gusA* DNA to replace both *brsM* and the counterselection cassettes all yielded numerous dark blue colonies, further confirming the efficacy of negative selection as well as the expected *gusA* allelic replacement of *brsM* (Fig. 3B). Importantly, despite the presence of some detectable background growth in the negative control reactions, the markerless *gusA* transformations all yielded few, if any, white colonies, indicating that nearly 100% of the transformants exhibited the expected Δ*brsM gusA*^*+*^ genotype (Fig. 3B).

**Fig. 3.**
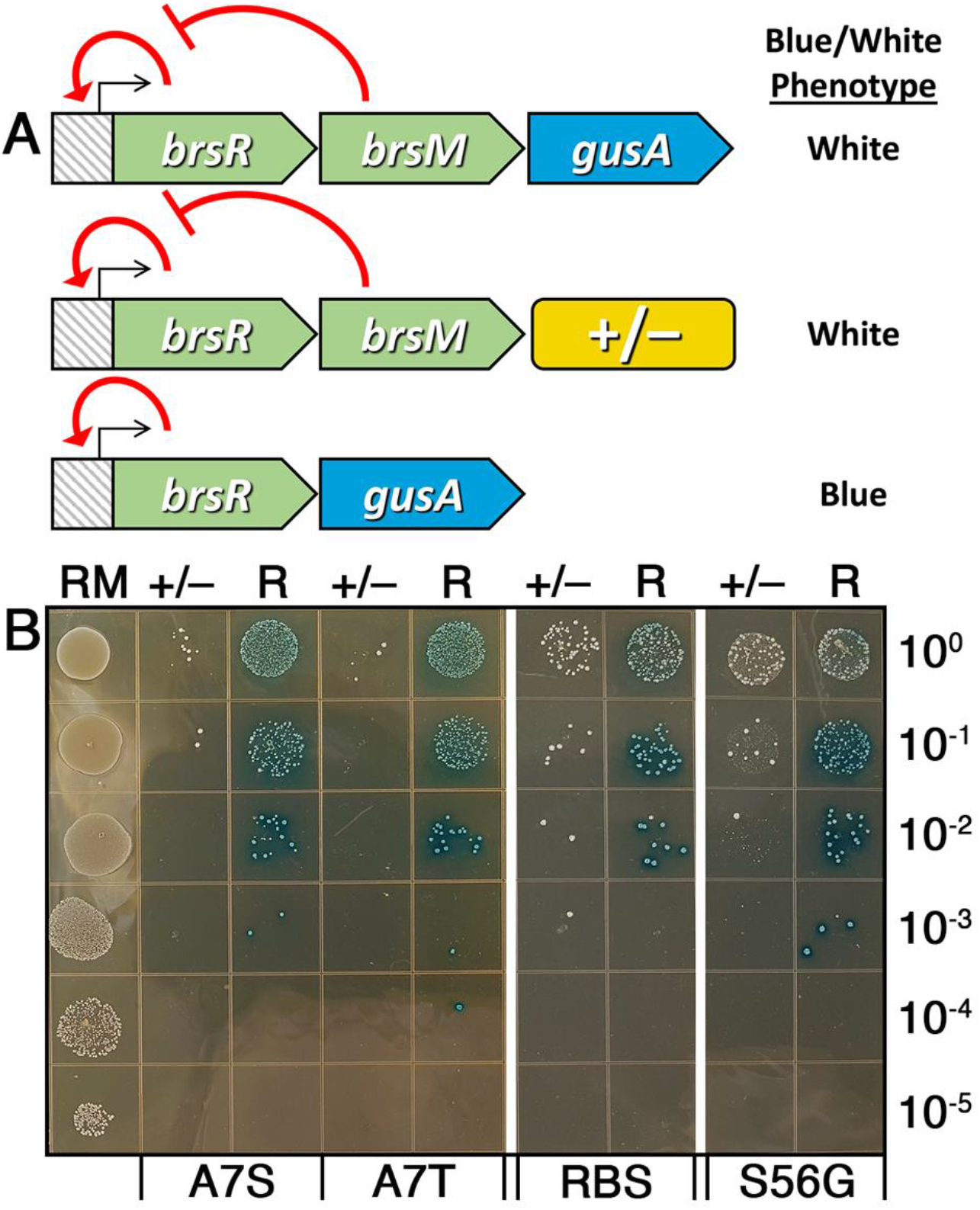
Allelic replacement with the *gusA* ORF. A) Illustration of the three genotypes encountered during construction of the Δ*brsM brsR-gusA* reporter strains. The top panel shows the previously constructed *brsRM-gusA* strain that was used as a template during construction of the markerless mutant strains. Since this strain has a wild-type *brsM* ORF, expression of the *brsRM* operon remains in its basal state due to the inhibitory function of BrsM towards the operon activator protein BrsR. Consequently, this strain will not exhibit detectable β-glucuronidase activity due to low *gusA* expression. The middle panel shows the genotype of strains transformed with the different counterselection cassettes (+/–), replacing the *gusA* ORF from the parent *brsRM-gusA* reporter strain. The bottom panel shows the genotype of the expected markerless reporter strains created by replacing both *brsM* and the different counterselection cassettes with the *gusA* ORF. After deleting *brsM*, inhibition of BrsR is relieved, resulting in potent autoactivation of the operon promoter and high levels of *gusA* expression. The markerless Δ*brsM brsR-gusA* reporter strains will exhibit a dark blue colony phenotype due to the large amount of β-glucuronidase activity produced from the reporters. B) The *brsRM-gusA* reporter strain (RM, left lane) was spotted onto xylose-supplemented agar plates in successive 10-fold dilutions. Each of the adjacent lanes to the right represent the transformation results of strains harboring counterselection cassettes transformed with either H_2_O (+/–) or DNA designed to replace *brsM* with the *gusA* ORF (R). The different counterselection cassettes are labeled as follows: Fst-sm A7S mutant (A7S), Fst-sm A7T mutant (A7T), *parE* RBS mutant (RBS), and ParE S56G mutant (S56G). The numbers on the right side of the image indicate the dilution factor of the cultures spotted onto the xylose plates.

### Comparison of the *fst-sm* and *parE* counterselection cassettes in multiple streptococci

As previously mentioned, one of our goals was to create a counterselection cassette exhibiting a broader host range than the previous *pheS-*based IFDC2 system. Since the *fst-sm* and *parE* cassettes were suitable for performing CIMM in *S. mutans* strain UA159, we next tested these same cassettes in three additional strains of *S. mutans* to determine whether they exhibit any evidence of strain-specificity. Both of the *fst-sm* cassettes performed quite similarly in each of the three *S. mutans* strains as previously observed with UA159. On xylose-supplemented agar plates, we detected 3 – 4 log reductions in total cell number, which corresponds to ≤0.1% background growth (Table 2). Unlike the *fst-sm* constructs, the results with the *parE* cassettes were mixed. The *parE* RBS point mutant version only exhibited noticeable negative selection in strain CL1 (Table 2), which is a serotype K clinical isolate (18). However, even in this strain, growth was partially inhibited on plates lacking xylose, which indicated that the basal expression of this toxin was still negatively affecting its growth. The ParE S56G mutant cassette performed better than the RBS mutant cassette, yielding xylose-inducible 3 – 4 log reductions in strains CL1 and JF243 with no obvious toxicity on the xylose-free plates (Table 2). Interestingly, strain UA140 was completely resistant to negative selection using both of the *parE* cassettes (Table 2). It is not yet clear why *parE* selection failed in UA140, but we suspect it is because this strain naturally harbors the pUA140 cryptic plasmid that encodes the ParE addiction module. From these results, we conclude that both of the *fst-sm* counterselection cassettes are likely to function well in most strains of *S. mutans*, while the ParE S56G cassette is slightly less universal, possibly due to the presence of the pUA140 plasmid in some strains.

**Table 2.** Quantification of inducible negative selection among streptococci.

| Species | Strain | Fst-sm A7S |  |  | Fst-sm A7T |  |  | ParE RBS |  |  | ParE S56G |  |  |
| --- | --- | --- | --- | --- | --- | --- | --- | --- | --- | --- | --- | --- | --- |
|  |  | – Xyl<br>CFU/ml | + Xyl<br>CFU/ml | Ratio | –Xyl<br>CFU/ml | +Xyl<br>CFU/ml | Ratio | –Xyl<br>CFU/ml | +Xyl<br>CFU/ml | Ratio | –Xyl<br>CFU/ml | +Xyl<br>CFU/ml | Ratio |
| *S. mutans* | UA140 | 4.3x10<sup>8</sup> | 1.2x10<sup>4</sup> | ≥10<sup>4</sup> | 3.4x10<sup>8</sup> | 1.5x10<sup>4</sup> | ≥10<sup>4</sup> | 6.4x10<sup>8</sup> | 5.2x10<sup>8</sup> | <2 | 7.0x10<sup>8</sup> | 6.5x10<sup>8</sup> | <2 |
| *S. mutans* | CL1 | 4.3x10<sup>8</sup> | 8.0x10<sup>4</sup> | ≥10<sup>3</sup> | 3.6x10<sup>8</sup> | 3.0x10<sup>4</sup> | ≥10<sup>4</sup> | 2.3x10<sup>6</sup> | 6.0x10<sup>3</sup> | ≥10<sup>2</sup> | 5.6x10<sup>8</sup> | 5.0x10<sup>4</sup> | ≥10<sup>4</sup> |
| *S. mutans* | JF243 | 7.8x10<sup>8</sup> | 7.0x10<sup>4</sup> | ≥10<sup>4</sup> | 7.5x10<sup>8</sup> | 6.0x10<sup>4</sup> | ≥10<sup>4</sup> | 4.8x10<sup>8</sup> | 4.5x10<sup>8</sup> | <2 | 7.6x10<sup>8</sup> | 1.0x10<sup>5</sup> | ≥10<sup>3</sup> |
| *S. gordonii* | DL1 | 5.8x10<sup>8</sup> | 4.8x10<sup>8</sup> | <2 | 7.5x10<sup>8</sup> | 6.0x10<sup>5</sup> | ≥10<sup>3</sup> | 6.6x10<sup>8</sup> | 8.0x10<sup>8</sup> | <1 | 8.8x10<sup>8</sup> | 6.5x10<sup>8</sup> | <2 |
| *S. sanguinis* | SK36 | 4.5x10<sup>8</sup> | 9.0x10<sup>6</sup> | >10 | 8.0x10<sup>8</sup> | 7.0x10<sup>5</sup> | ≥10<sup>3</sup> | 7.6x10<sup>8</sup> | 5.0x10<sup>5</sup> | ≥10<sup>4</sup> | 6.8x10<sup>8</sup> | 7.2x10<sup>8</sup> | <1 |

Since we had previously demonstrated the utility of the Xyl-S system for xylose-inducible gene expression in both *S. gordonii* and *S. sanguinis* (17), we were also curious to determine whether the xylose-inducible *fst-sm* and *parE* counterselection cassettes would function in these organisms. We introduced the four *fst-sm* and *parE* counterselection cassettes into *S. gordonii* strain DL1 and *S. sanguinis* strain SK36, which are among the most commonly utilized strains for genetic studies of both species. Only the *fst-sm* (A7T) cassette functioned in both organisms, yielding 3-log reductions in the presence of xylose (Table 2). Surprisingly, the RBS mutant *parE* cassette functioned even better than *fst-sm* (A7T) in *S. sanguinis*, but it completely failed to inhibit *S. gordonii* (Table 2). The *fst-sm* (A7S) and *parE* (S56G) cassettes did not function in either *S. gordonii* or *S. sanguinis*. Overall, the negative selection results indicated that the *fst-sm* (A7T) cassette is likely to be the most broadly useful counterselection cassette for streptococci, although the *fst-sm* (A7S) and the *parE* cassettes may also function quite well for certain species or strains. To further confirm the utility of the *fst-sm* (A7T) counterselection cassette in *S. gordonii* and *S. sanguinis*, we performed CIMM with this cassette to markerlessly insert the green renilla luciferase ORF (*renG*) immediately downstream of the pyruvate oxidase-encoding gene *spxB* in both species (Fig. 4A). Since the pyruvate oxidase enzyme is responsible for generating the majority of the H_2_O_2_ excreted by these two streptococci (19, 23), the *spxB*^*–*^ phenotype is readily observable on Prussian blue plates, as this dye reacts with H_2_O_2_ to produce a blue precipitate. As shown in figure 4B, both *S. gordonii* and *S. sanguinis* produced much less blue precipitate when *spxB* was replaced with the *fst-sm* (A7T) counterselection cassette. However, wild-type levels of H_2_O_2_ were once again detectable when *spxB* was markerlessly inserted back to its original locus together with a downstream *renG* ORF (Fig. 4B). We also measured luciferase activity from both of the markerless luciferase reporter strains and observed ∼4-log increased reporter activity over the background, unlike the parent *spxB* deletion strains, which lacked the *renG* ORF (Fig. 3C). Thus, we conclude that the *fst-sm* (A7T) counterselection cassette (henceforth referred to as IFDC4) is suitable for both markerless deletions and insertions in *S. gordonii* and *S. sanguinis*.

**Fig. 4.**
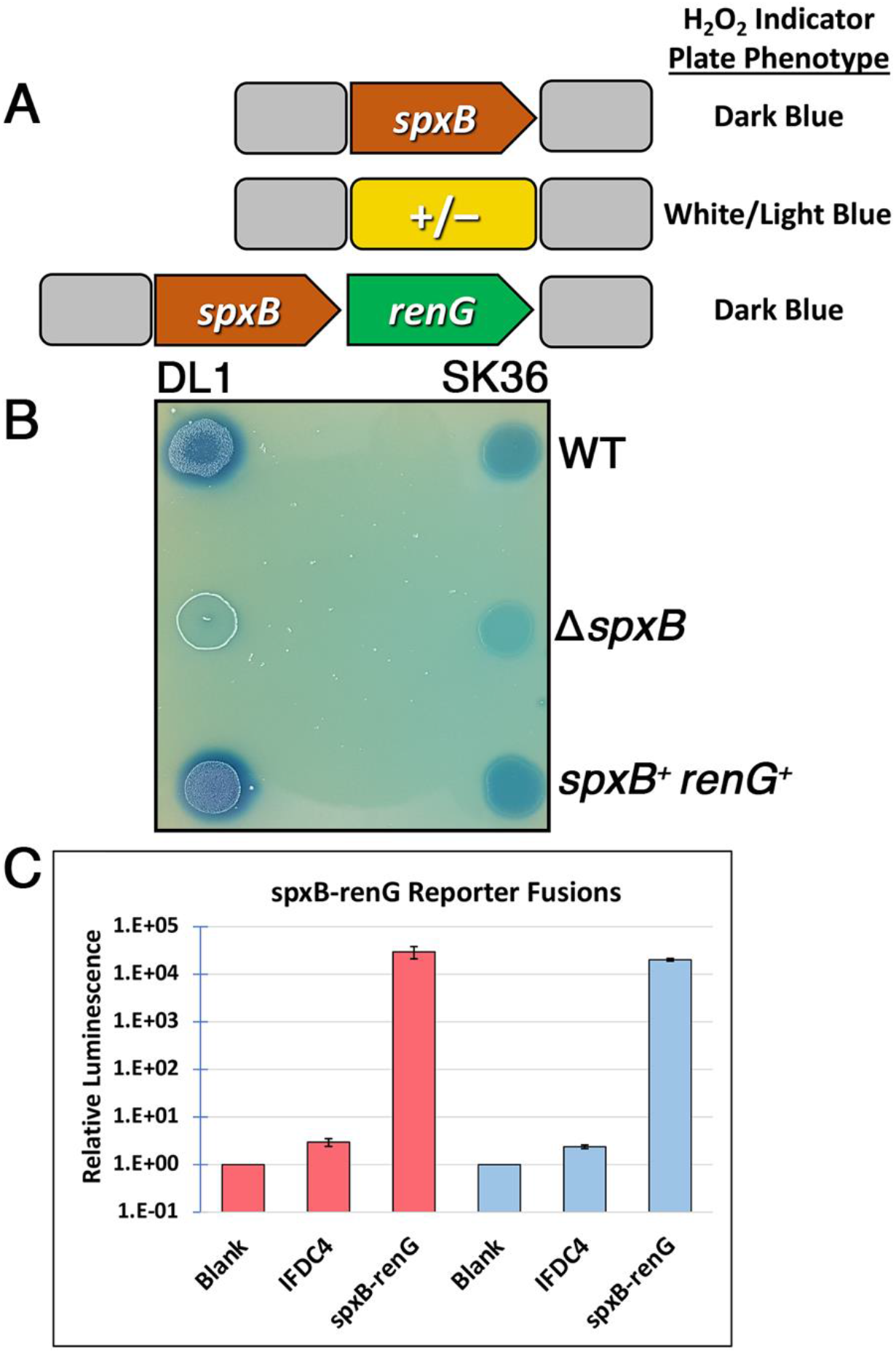
Markerless gene deletions and insertions in *S. gordonii* and *S. sanguinis*. A) Illustration of the three genotypes encountered during construction of the markerless *spxB-renG* reporter strains. The top panel shows the wild-type *spxB* locus of both *S. gordonii* and *S. sanguinis*. Both wild-type strains yield a blue precipitate on Prussian blue agar plates due to the H_2_O_2_ produced primarily from the *spxB-*encoded enzyme pyruvate oxidase. The middle panel shows an allelic replacement of *spxB* with the counterselection cassettes (+/–), resulting in a major reduction in H_2_O_2_ production. A subsequent transformation of these strains with *spxB-renG* DNA replaces IFDC4 and restores *spxB* to its original locus along with a transcriptional fusion to the *renG* ORF. The markerless *spxB-renG* reporter strains should produce similar levels of H_2_O_2_ as the original wild-type strains. B) Representative strains of the wild-type (WT), *spxB* mutant (Δ*spxB*), and *spxB-renG* reporter (*spxB*^*+*^ *renG*^*+*^) of both *S. gordonii* (DL1) and *S. sanguinis* (SK36) were spotted onto Prussian blue agar plates to observe the H_2_O_2_ production phenotypes of each. C) Luciferase activity was compared between the Δ*spxB* strains harboring the counterselection cassettes (IFDC4) vs. the markerless *spxB-renG* reporter strains (spxB-renG). Results from the *S. gordonii* strains are shown in red, while *S. sanguinis* results are shown in blue. Luciferase data are presented relative to the cell-free background luciferase values, which were arbitrarily assigned a value of 1. Luciferase data are derived from five independent clones of each strain, which were averaged and presented together with their corresponding standard deviations.

### Markerless point mutagenesis

As previously described, one potential advantage of counterselection-based markerless mutagenesis is that it supports the creation of targeted point mutations due to scarless excision of the counterselection cassettes. Therefore, as a final confirmation of IFDC4 utility, we designed CIMM constructs to create nonsense point mutations within the *galK* genes of *S. mutans, S. gordonii*, and *S. sanguinis*. We chose to mutate *galK* because this gene is both nonessential in all three species and should confer resistance to the toxic effects of the galactose analog deoxygalactose, yielding an easily observable growth phenotype. As shown in figure 5A, deoxygalactose resistance was indeed created after markerlessly introducing an ochre (CAA → TAA) nonsense point mutation into codon 8 of the *S. mutans* UA159 *galK* gene. We repeated the same experiment using *S. gordonii* DL1 and were able to successfully introduce a similar ochre (CAA → TAA) nonsense mutation into codon 9 of its *galK* gene (Fig. 5B). However, we were surprised to discover that wild-type DL1 is naturally resistant to the toxic effects of deoxygalactose. Therefore, unlike *S. mutans*, we were unable to observe any difference in growth between the wild-type and *galK* point mutant strains. For unknown reasons, we could not achieve any discernable negative selection when trying to mutate the *galK* gene of *S. sanguinis* SK36, despite the efficacy of negative selection with our other SK36 mutant constructs using the same IFDC4 cassette (Table 2 and Fig. 4B – C). We even repeated the experiment using the ParE RBS mutant counterselection cassette, which exhibited even stronger negative selection in SK36 compared to IFDC4 (Table 2), yet we still observed the same problem. This issue appeared unique to the *galK* locus, as negative selection was quite stringent for our other SK36 constructs (Table 2 and Fig. 4B – C), and we have also recently used IFDC4 to engineer additional markerless mutations in SK36 for our other ongoing research. It is unclear why the SK36 *galK* locus is specifically problematic, but based upon our results, it would appear that this mutation affects the utility of xylose induction. However, additional studies would be required to determine whether this is indeed the cause. Regardless, our success with both *S. mutans* and *S. gordonii* suggests that point mutagenesis would be unlikely to fail in most instances in SK36.

**Fig. 5.**
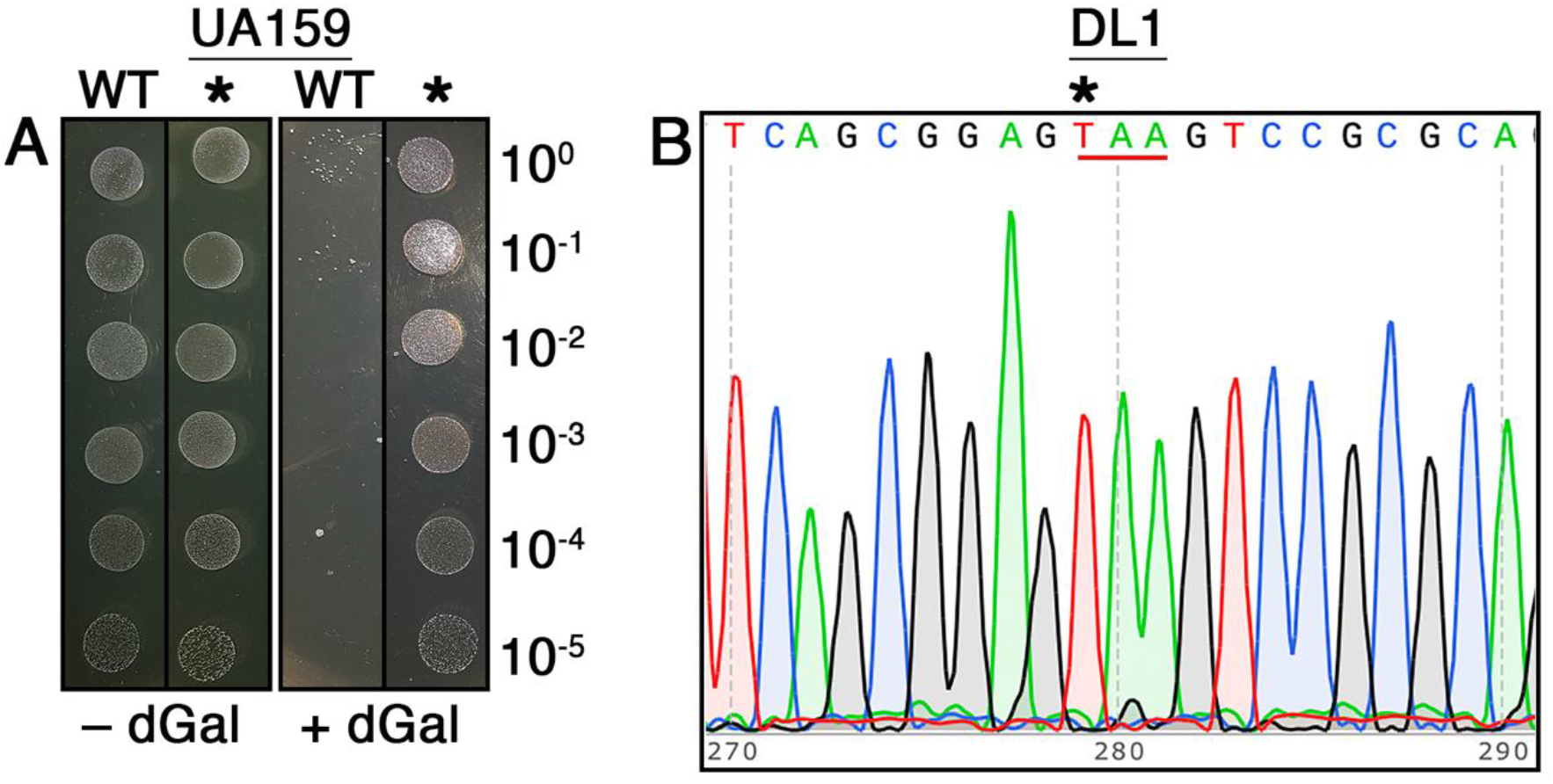
Introduction of markerless *galK* point mutations. A) IFDC4 was used to engineer a markerless nonsense point mutation into the *galK* gene of *S. mutans*. After sequencing to confirm the presence of the expected stop codon, the mutant strain (*) was spotted adjacent to the parent wild-type strain (WT) in successive 10-fold dilutions on agar plates ± deoxygalactose (dGal). The numbers on the right side of the image indicate the dilution factor of the cultures spotted onto the agar plates. B) Sequence results of the *S. gordonii galK* gene following IFDC4 point mutagenesis. The engineered stop codon is underlined, while the specific C→T mutation site is marked with an asterisk.

## Discussion

In the current study, we describe a new approach to perform CIMM in streptococci. This system was developed to address a couple of the limitations encountered when using the previous *pheS-*based IFDC2 counterselection cassette, mainly its requirement for a toxic substrate during negative selection (i.e. 4-CP) and its narrow host range. Here, we employed the Xyl-S induction system to control the expression of a toxic gene product as a negative selection mechanism. We have previously used the Xyl-S system to introduce controllable gene expression in a number of *in vitro* and *in vivo* studies (16, 17), so this expression system has proven utility in multiple streptococci with no evidence of xylose toxicity. Furthermore, our previous studies had already suggested that the Xyl-S system would be suitable for engineering conditional lethality (17), which we repurposed here as a negative selection mechanism for CIMM. As shown in Table 2, negative selection was highly efficient with exceptionally low background, resulting in an extremely high percentage of markerless mutants with the expected genotypes (Fig. 3B). Unlike the IFDC2 cassette, IFDC4 also allowed us to employ a single cassette to engineer a variety of different types of markerless mutations in *S. mutans, S. gordonii*, and *S. sanguinis*, thus confirming its broader host range. For unknown reasons, we were unable to induce negative selection when mutating the *galK* gene of *S. sanguinis*. This problem seemed specific to the *S. sanguinis galK* locus, as we did not encounter similar issues with either of the *S. mutans* or *S. gordonii galK* mutations (Fig. 5A – B), nor did we experience issues with negative selection in other *S. sanguinis* loci (Table 2 and Fig. 4B – C). We tried a variety of different construct designs to create the *galK* mutant, but all failed to exhibit any evidence of xylose-inducible negative selection. Therefore, we suspect that xylose is either modified or exhibits defective transport following mutagenesis of the *S. sanguinis galK* locus.

Our results provide a general strategy for negative selection that should be adaptable for use in other organisms. Most bacteria encode endogenous toxin-antitoxin modules on their chromosomes, while many others also have particular strains that naturally host cryptic plasmids, which are highly likely to contain addiction modules (24, 25). Thus, there is a strong reservoir of potential toxins that could be exploited for use in most bacteria. This approach also requires a reliable regulated gene expression system, which may be a limiting factor for certain organisms, especially those with minimal genetic tools available. From our experience, successful toxin-based negative selection requires both low basal uninduced expression of the toxin gene as well as strong inducibility (i.e. wide dynamic range of expression). However, the specific expression characteristics required to create an efficacious inducible negative selection system are likely to vary widely. Thus, the challenge is to find an appropriate match between the available expression system(s) and the toxicity of a particular toxic gene product. For example, our results suggest that ParE is likely to be a more potent toxin in *S. mutans* compared to Fst-sm, as *fst-sm* could be stably expressed from the Xyl-S expression system after acquiring a single point mutation within the *xylR* gene, whereas the *parE* constructs required the same *xylR* point mutation as well as additional compensatory *parE* mutations (Fig. 2C). Thus, when developing a new toxin-based negative selection system, it is advisable to screen a variety of candidate toxins to increase the likelihood of identifying the appropriate gene to pair with an available expression system. In our case, none of the toxin genes we selected were immediately ideal for our Xyl-S expression system. However, the *fst-sm* construct was apparently a sufficiently close match that it allowed us to isolate spontaneous compensatory xylose repressor mutations, resulting in a mutant counterselection cassette with lower basal uninduced toxicity, but stringent xylose-inducible negative selection. It seems unlikely that we would have identified the appropriate compensatory *parE* mutations if the strain did not already contain the XylR A7S mutation within the Xyl-S induction system, which apparently reduced its leakiness. This may also explain why we did not isolate spontaneous compensatory mutations for the other toxins in our initial screen of candidates, as these were constructed without the XylR A7S or A7T mutations. Presumably, the mutants we did isolate with these other toxins all contained inactivating mutations because none of the isolates exhibited any evidence of xylose-inducible negative selection (Fig. 2A). Since we had already observed effective negative selection from the *fst-sm* and *parE* constructs, we did not perform additional tests to determine whether the other toxin genes might function in combination with the XylR A7S mutation. However, it is certainly conceivable that at least some of them would, perhaps requiring additional compensatory mutations similar to the *parE* construct. As our results demonstrate, it is not essential for a given gene induction system to perfectly pair with a toxin gene, provided the initial basal uninduced toxicity of the construct is moderate enough that it gives the cell a chance to develop the appropriate compensatory mutations. From there, one can screen the isolates to identify those that may be suitable for stable negative selection.

